# Identification of BLM’s Aggregation-Prone Nature and Its Similarity to Intrinsically Disordered Proteins: An In-silico Study

**DOI:** 10.1101/2024.07.16.603833

**Authors:** Sourav Sharma, Suresh Singh Yadav, Rohini Ravindran Nair

## Abstract

BLM helicase is a member of the RecQ family of DNA helicases, enzymes that play a critical role in maintaining genome stability. The BLM protein is named after Bloom syndrome (BS), a rare genetic disorder caused by mutations in the helicase domain of BLM gene. BLM role has mostly been studied in DNA repair, replication, and recombination; however, a recent study has highlighted the RNA binding capability of the BLM protein. Several proteins, including TDP43, Tau, and alpha-synuclein, are known to play key roles in neurodegenerative diseases. These proteins possess residues prone to parallel aggregation and fibril formation, the key contributor to neurodegenerative disease development. We utilized various in-silico tools like PASTA 2.0, SIFT, PolyPhen 2, and PhD SNP, SNP&GO, Meta-SNP, and SNAP, etc., to identify the molecular signature of the BLM’s protein. Our in-silico analysis suggests that the BLM’s HRDC domain has ability of parallel aggregation and fibril formation. Moreover, structural similarity with proteins like α-synuclein, TDP-43, and Tau and its interactions with PARP1 suggests it may have a role in neurodegenerative diseases. Furthermore, we identified deleterious mutations in the HRDC domain of BLM protein that may compromise its stability and alter its function. Hence, these findings suggest that in addition to BLM’s well-known functions, the protein may have ability to form parallel aggregation and fibril formation and its role in neurodegenerative disease need to further be explored.

**Author Summary:** This paper higlights the importance of considering the structural and physical characteristics of the BLM protein, which may have been previously overlooked in the context of disease pathology. Through in-silico analyses, we identified that the BLM protein possesses residues within the HRDC domain that are prone to parallel aggregation and fibril formation. Moreover, the structural similarity of BLM to proteins such as α-synuclein, TDP-43, and Tau indicates a potential role for BLM in neurodegenerative processes. Additionally, we identified deleterious mutations within the HRDC domain that could compromise the stability of the protein. Furthermore, our findings suggest that the association between BLM and PARP1 may involve a regulatory mechanism that could significantly influence BLM’s function.

## Introduction

BLM’s is well-known for its role in repairing DNA double-strand breaks through homologous recombination. It plays a key part in DNA end resection, D-loop formation, Rad51 filament formation, branch migration, and the dissolution of Holliday junctions [1]. Additionally, BLM efficiently unwinds DNA secondary structures, with a particularly strong ability to unwind G-quadruplexes, making it one of the most effective human DNA helicases in this regard. Recent research has revealed that BLM also binds to RNA, showing a preference for binding to RNA G-quadruplexes (G4s) [2]. RNA binding proteins (RBPs) are known to have role in neurodegenerative disease, and they also share a binding preference for G-quadruplexes (G4s) [3].

Neurodegenerative diseases are associated to a group of proteins referred as intrinsically disordered proteins (IDPs) [4]. IDPs are a normal class of proteins, but unlike most proteins that fold into specific three-dimensional structures, IDPs or proteins with intrinsically disordered regions (IDRs) do not have a fixed structure under physiological conditions. This lack of a defined structure allows them to be highly flexible and versatile, enabling them to interact with multiple different partners and play key roles in various biological processes. Alterations in the cellular environment or mutations within IDPs can interfere with their normal functions, leading to protein misfolding and the formation of aggregates or fibrils [5, 6]. Studies have reported the association of IDPs/IDRs proteins such as TAR DNA-binding protein (TDP-43) in amyotrophic lateral sclerosis (ALS), α-synuclein (α-syn) in Parkinson’s disease (PD), and amyloid beta (Aβ) and tau for Alzheimer’s disease (AD) [7].

Protein posttranslational modifications (PTMs) of IDPs/IDRs also determine the formation of pathologic aggregation. One of the known PTMs involved in neurodegeneration is poly (ADP-ribosylation) (PARylation). A family of PARP enzymes performs PARylation that attaches multiple NAD-derived ADP-ribose (ADPr) units to the acceptor protein [8]. The number of cellular processes such as gene expression, cell death pathway, DNA repair, mitochondrial biogenesis, and neuroinflammation, are regulated by PARylation and PAR-binding [9-15]. PARP1 activity is known to be elevated in the brains of individuals with neurodegenerative diseases, including Alzheimer’s disease. Several proteins involved in Alzheimer’s disease (AD) and amyotrophic lateral sclerosis (ALS), such as α-synuclein, TDP-43, and hnRNP A1, are regulated by PARylation and PAR-binding. These modifications influence the phase separation properties and aggregation tendencies of these proteins [16-18].

In this study, we utilized in-silico tools to explore the functional aspects of BLM, particularly its potential to form aggregation, which may contribute to the pathology of neurodegenerative diseases. Using the PASTA 2.0 tool, we identified amino acid residues within the HRDC domain of the BLM protein that are likely to undergo parallel aggregation and fibril formation. Notably, our findings reveal significant similarities between BLM and intrinsically disordered proteins (IDPs) implicated in neurodegenerative diseases. Moreover, the mutation identified in the HRDC domain of BLM impacts its stability, further suggesting it may have a potential role in disease pathology. Additionally, our study uncovered critical associations between BLM and PARP1. Based on these insights, we suggest that the HRDC domain of BLM possesses aggregation and fibril formation capabilities, which may contribute to the pathophysiology of neurodegenerative diseases.

## Results

### BLM’s Expression Across Cell Types and Brain Regions

The human BLM belongs to the RecQ helicase family of proteins consisting of 1417 amino acids encoded by the BLM gene on chromosome 15 (UniProtKB ID: P54132) and has a molecular weight of approximately 159 kDa **(Fig 1A)**. This protein has a conserved ATP-dependent DNA helicase domain ranging from 657-865 amino acids. BLM also has RecQ C-terminal (RQC; 1072-1194) and helicase-and-ribonuclease D-C-terminal (HRDC; 1212-1292) domains. HRDC domain has single-stranded DNA (ssDNA) and the DNA Holliday junction binding residues between 1227-1244 amino acids. The unstructured regions of the protein include the N-terminus (residues 1–657), which is crucial for interaction and oligomerization with partner proteins, and the C-terminus (residues 1293–1417), which has nuclear localization signals and is crucial for the nuclear function of BLM.

**Figure 1:**
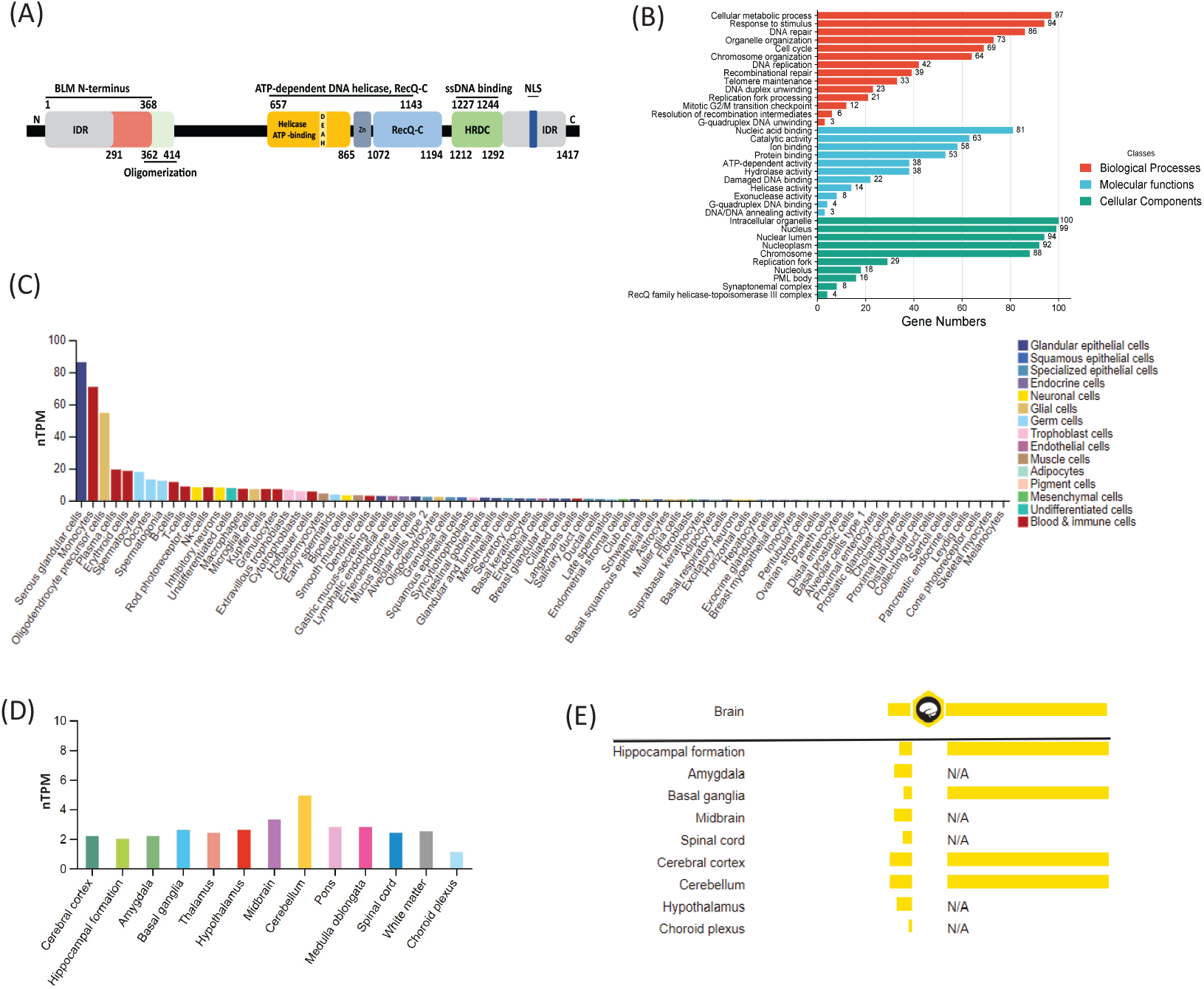
**(A)** Representation of BLM gene showing IDR (Intrinsically Disordered Regions), and Evolutionary conserved regions like Helicase-ATP binding domain, RecQ-C, and HRDC domain. **B)** Gene ontology (analysis) showing selected GO terms in Biological Processes, Molecular Functions, and Cellular Components for BLM interacting genes according to SR plot. **C)** A summary of normalized single-cell RNA nTPM for BLM from all single-cell types. Color coding is based on cell type groups, each consisting of cell types with functional features in common. **D)** Normalized RNA expression levels (nTPM) of BLM were shown for the 13 brain regions. Color coding is based on brain region and the bar shows the highest expression among the subregions included. **E)** Bar Graph representation of RNA and Protein expression for Brain tissue according to The Human Protein Atlas.

Gene Ontology (GO) analyses [16, 19, 20] indicated that BLM participates in diverse regulatory complexes in the cytoplasm and nucleus **Fig 1B**. Mostly its function has been reported on cellular metabolic processes such as DNA repair, and replication. Analysis of its expression pattern revealed its highest expression in serous glandular cells (**Fig 1C**). Oligodendrocyte precursor cells were also among the cells that had the third highest level of BLM gene expression after monocytes. On comparing the expression level in different regions of the brain its expression was found to be highest in the cerebellum. The expression of BLM at the protein level has not been identified in some regions of the brain even after its expression at the transcript level **Fig 1D and E**. Hence from the above finding we can suggest that BLM is expressed in neuronal cells and it may have a role in neuronal development and its function.

### BLM’s Parallel Aggregation and Fibril Formation Ability

On computing the physical and chemical parameters of BLM using ProtParam it was classified as an unstable protein with an instability index of 44.57 (smaller than 40 is considered as stable protein). The aliphatic index of BLM is 73.83, implying that the protein is thermostable. The GRAVY score calculated by the server was -0.602, indicating that the protein is hydrophilic. The estimated half-life is 30 hours (mammalian reticulocytes, in vitro), >20 hours (yeast, in vivo), >10 hours (Escherichia coli, in vivo) in the respective cell types. PASTA 2.0 predicted that a significant region of BLM contains 38.96% alpha-helical structure, 12.07% forms the beta-strands secondary structure, 48.98% forms coils and about 23.42% of the region remains disordered. According to PASTA 2.0, the BLM protein’s N-terminal residues 1-21, 39-42, 72-75, 92-107, 142-175, 183-232, 247-288, 290, 291, 313, and 323–339 was found to be disordered, while residues 1224–1250 show a probability of parallel aggregation and fibril formation (**Fig 2A**). Additionally, the C-terminal residues 1277-1417 were found to be disordered. This suggests that BLM has intrinsically disordered regions and has the ability of parallel aggregation and fibril formation.

**Figure 2:**
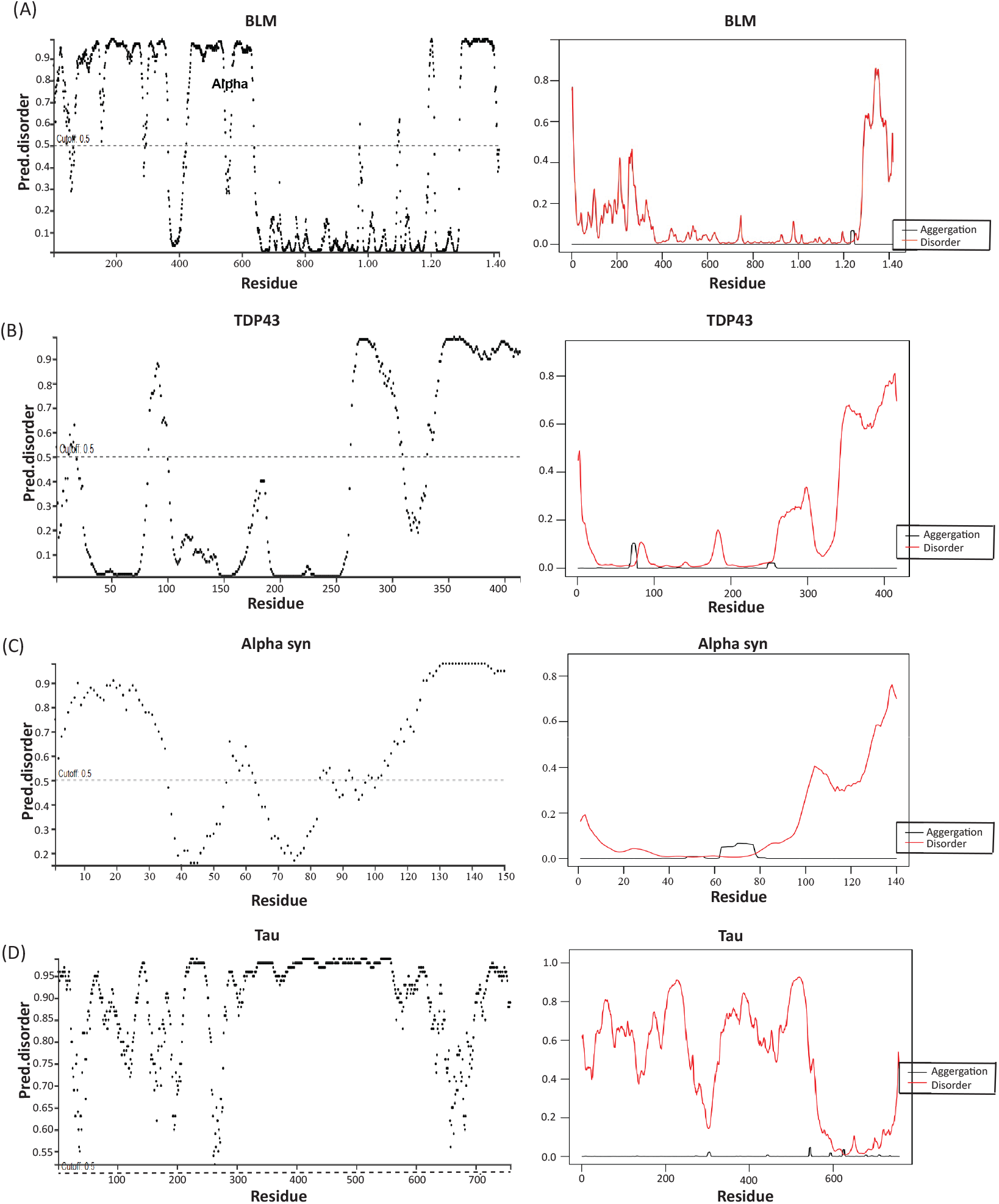
Depiction of Disordered and parallel aggregation formation plots: **A)** BLM (UniProt ID: P54132), **B)** TDP43 (UniProt ID: Q13148), **C)** α-Synuclein (UniProt ID: P37840), and **D)** Tau (UniProt ID: P10636) using DISOPRED3 and PASTA 2.0.

### BLM’s Similarity with TDP43, Tau, and A-Synuclein

Similar to TDP-43, α-syn, Aβ, and Tau, BLM (Bloom syndrome protein), also carries intrinsically disordered regions. By comparing the physical and chemical parameters of BLM with TDP43, Tau, and α-synuclein, we demonstrated that all are unstable with an instability index of 44.57, 56.76, and 56.02 respectively except α-synuclein which has an instability index of 25.47. Also, BLM, TDP43, Tau, and α-synuclein are all hydrophilic proteins with GRAVY scores of -0.602, -0.535, -0.882, and -0.403 respectively.

We used PSIPRED and DISOPRED3 webservers [21] to predict the secondary structure and intrinsically disordered regions of BLM, TDP43, Tau, and α-synuclein. The prediction revealed the presence of structural domains like α-helix and beta-strands strands in the ordered region of these proteins. Interestingly we noticed all had a disordered C-terminus region. Beta-strands were identified as regions in BLM protein with a threshold <0.5 to form parallel aggregation using the PASTA 2.0 tool (**Fig 2)**. Hence the ability of BLM protein to form parallel aggregation is similar to TDP43, Tau, and α-synuclein which are commonly involved in neurodegenerative diseases like amyotrophic lateral sclerosis (ALS), Parkinson’s disease (PD), and Alzheimer’s disease (AD) **(Table 1**).

**Table 1.**
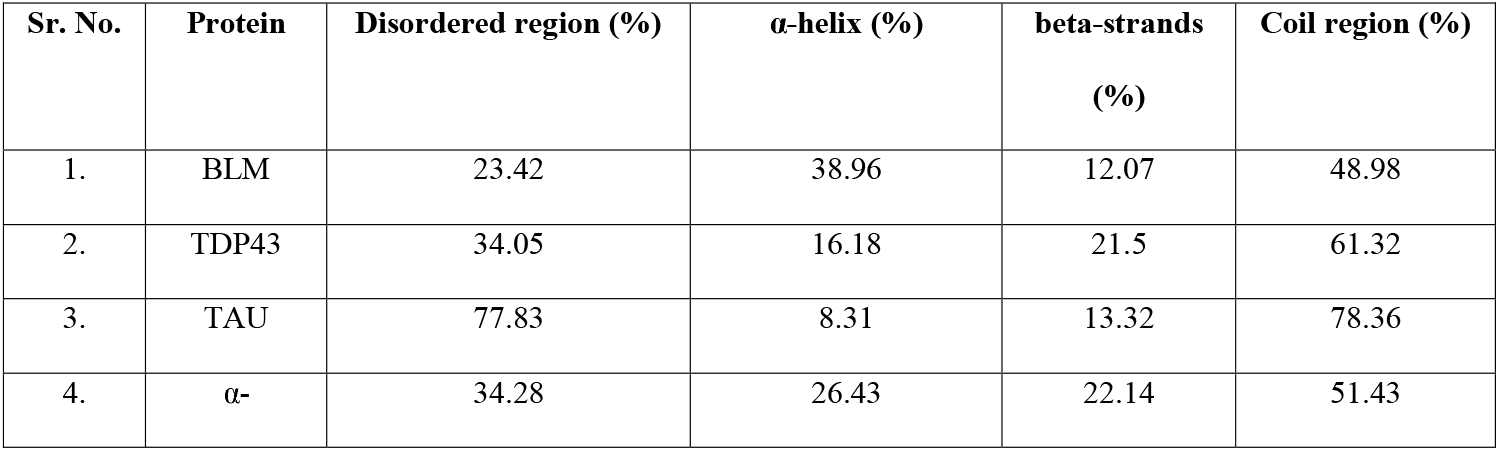
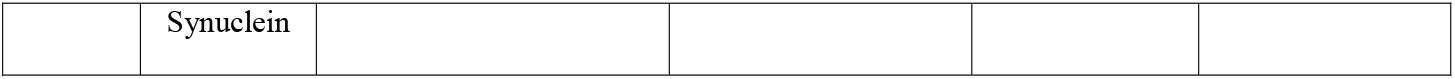
Structural Composition and Secondary Structure Elements of BLM, TDP43, Tau, and α-Synuclein Proteins.

| Sr. No. | Protein | Disordered region (%) | α-helix (%) | beta-strands (%) | Coil region (%) |
| --- | --- | --- | --- | --- | --- |
| 1. | BLM | 23.42 | 38.96 | 12.07 | 48.98 |
| 2. | TDP43 | 34.05 | 16.18 | 21.5 | 61.32 |
| 3. | TAU | 77.83 | 8.31 | 13.32 | 78.36 |
| 4. | α- | 34.28 | 26.43 | 22.14 | 51.43 |

|  |  |
| --- | --- |
|  | Synuclein |

### Identification Of Deleterious and Damaging nsSNPs in BLM

Mutations in BLM can affect the stability of protein hence to identify the most potentially harmful nsSNPs in BLM, we used the ENSEMBL browser displaying 44201 genetic variations (Transcript: ENST00000355112.8 (BLM-201) - Variant table - Homo_sapiens - Ensembl genome browser 112). Furthermore, these SNPs were further validated by the NCBI Variation Viewer and dbSNP database (BLM: Chr15 -Variation Viewer (nih.gov). Using both of these tools we identified 40919 single nucleotide variants of the BLM gene. Using the genomic coordinates of the “NM_000057.4” transcript of the BLM helicase isoform 1, we identified 2647 SNPs in the coding region sequence. These included missense variants (non-synonymous SNPs), 123 stop-gained variants, and 2 start-lost nsSNPs, as illustrated in **Fig 3**. The remaining nsSNPs were filtered based on the standard scoring of various sequence-based consensus tools: SIFT (score <0.05) [22]; PolyPhen-2 (score>0.95 - probably harmful, score>0.5 – possibly damaging) [23]; Mutation Assessor (score>0.5); Meta-SNP, PhD-SNP, SNPs & Go (score>0.5) [24]. The filtered nsSNPs using the above tools were further validated using PredictSNP [25] & SNAP [26] to determine the accuracy of deleterious mutations.

**Figure 3:**
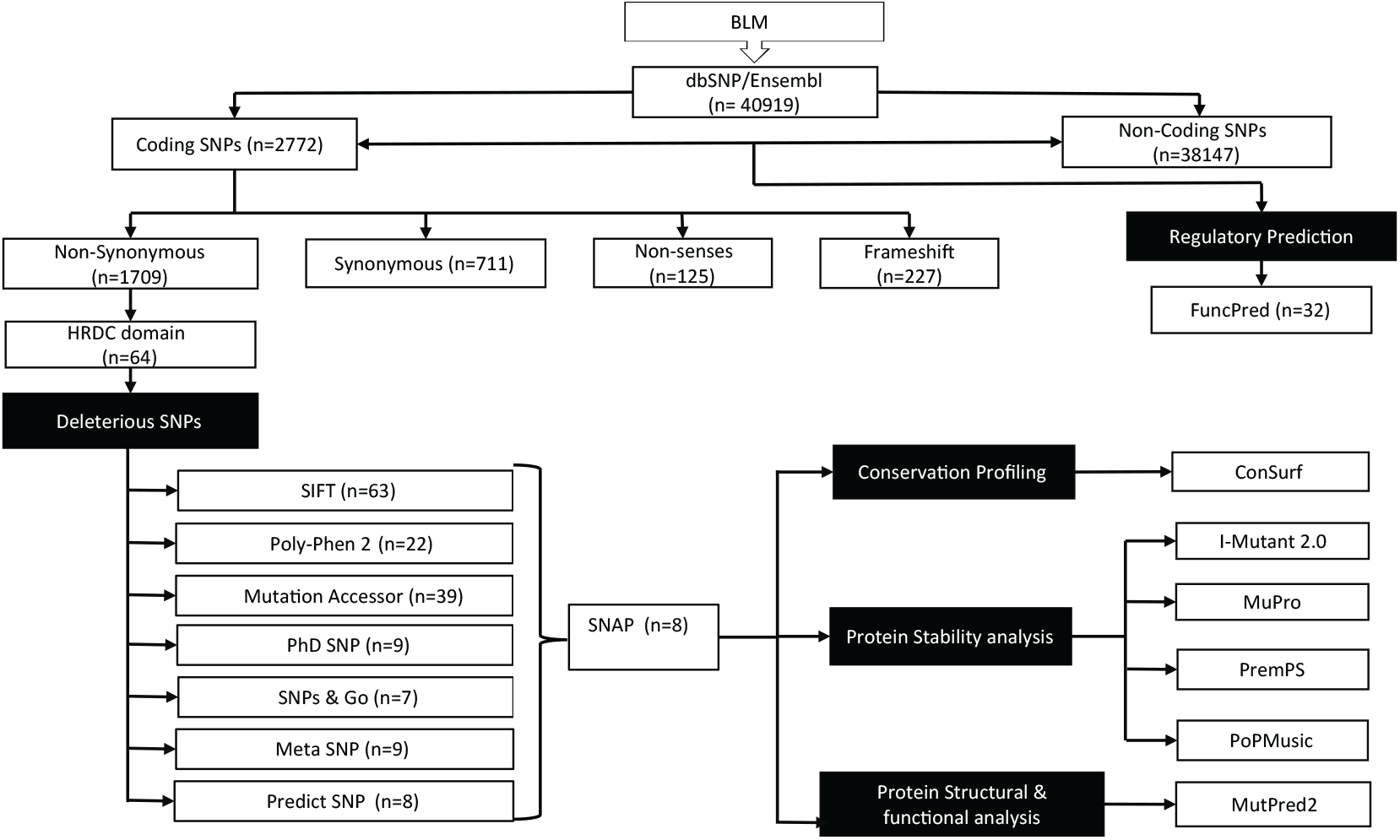
A schematic representation of nsSNPs selection and shortlisting procedure by different tools.

We further examined the highly conserved HRDC (helicase and RNase D C-terminal) domain that was found to form parallel aggregations (**Fig 2A**). A total of 64 nsSNPs was found in the HRDC domain and based on the score predicted by the above sequence-based tools, 8 nsSNPS (**Table 2**) (H1236Y, F1238L, L1253S, T1267I, E1268V, E1272G, V1278M, S1286) was classified as predominantly deleterious. Various studies have shown that most disease-related missense mutations alter the stability of proteins. We analyzed these nsSNPs for amino acid substitutions and their impact on the stability of mutant BLM protein using I-Mutant 2.0 (http://folding.biofold.org/i-mutant/i-mutant2.0.html) [27], MuPro (https://mupro.proteomics.ics.uci.edu) [28], PremPS (https://lilab.jysw.suda.edu.cn/research/PremPS/) [29] and PoPMuSiC v3.1 2.1 (http://babylone.ulb.ac.be/PoPMuSiCv3.1) [30].

**Table 2.**
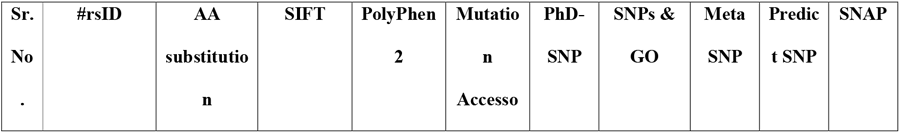
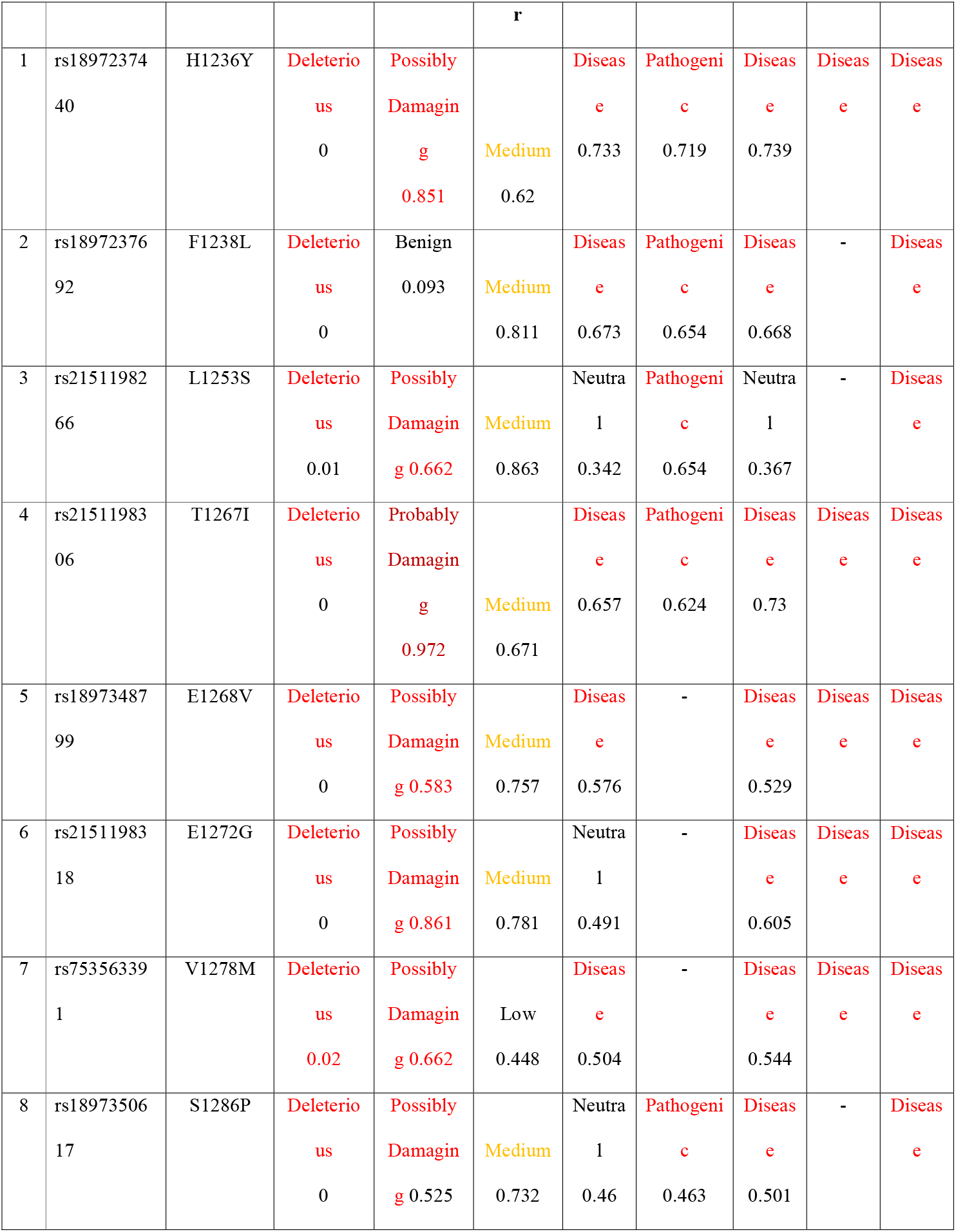
List of deleterious missense SNPs in the HRDC domain of BLM gene using different sequence-based tools.

| Sr. No. | #rsID | AA substitution | SIFT | PolyPhen 2 | Mutation Accessor | PhD-SNP | SNPs & GO | Meta SNP | Predict SNP | SNAP |
| --- | --- | --- | --- | --- | --- | --- | --- | --- | --- | --- |

|  |  |  |  |  | r |  |  |  |  |  |
| --- | --- | --- | --- | --- | --- | --- | --- | --- | --- | --- |
| 1 | rs18972374<br>40 | H1236Y | Deleterio<br>us<br>0 | Possibly<br>Damagin<br>g<br>0.851 | Medium<br>0.62 | Diseas<br>e<br>0.733 | Pathogeni<br>c<br>0.719 | Diseas<br>e<br>0.739 | Diseas<br>e | Diseas<br>e |
| 2 | rs18972376<br>92 | F1238L | Deleterio<br>us<br>0 | Benign<br>0.093 | Medium<br>0.811 | Diseas<br>e<br>0.673 | Pathogeni<br>c<br>0.654 | Diseas<br>e<br>0.668 | - | Diseas<br>e |
| 3 | rs21511982<br>66 | L1253S | Deleterio<br>us<br>0.01 | Possibly<br>Damagin<br>g<br>0.662 | Medium<br>0.863 | Neutra<br>l<br>0.342 | Pathogeni<br>c<br>0.654 | Neutra<br>l<br>0.367 | - | Diseas<br>e |
| 4 | rs21511983<br>06 | T1267I | Deleterio<br>us<br>0 | Probably<br>Damagin<br>g<br>0.972 | Medium<br>0.671 | Diseas<br>e<br>0.657 | Pathogeni<br>c<br>0.624 | Diseas<br>e<br>0.73 | Diseas<br>e | Diseas<br>e |
| 5 | rs18973487<br>99 | E1268V | Deleterio<br>us<br>0 | Possibly<br>Damagin<br>g<br>0.583 | Medium<br>0.757 | Diseas<br>e<br>0.576 | - | Diseas<br>e<br>0.529 | Diseas<br>e | Diseas<br>e |
| 6 | rs21511983<br>18 | E1272G | Deleterio<br>us<br>0 | Possibly<br>Damagin<br>g<br>0.861 | Medium<br>0.781 | Neutra<br>l<br>0.491 | - | Diseas<br>e<br>0.605 | Diseas<br>e | Diseas<br>e |
| 7 | rs75356339<br>1 | V1278M | Deleterio<br>us<br>0.02 | Possibly<br>Damagin<br>g<br>0.662 | Low<br>0.448 | Diseas<br>e<br>0.504 | - | Diseas<br>e<br>0.544 | Diseas<br>e | Diseas<br>e |
| 8 | rs18973506<br>17 | S1286P | Deleterio<br>us<br>0 | Possibly<br>Damagin<br>g<br>0.525 | Medium<br>0.732 | Neutra<br>l<br>0.46 | Pathogeni<br>c<br>0.463 | Diseas<br>e<br>0.501 | - | Diseas<br>e |

Currently, the whole protein 3D structure of the BLM gene is not available, hence, partial protein structures, such as PDB id 4O3M with a length of 640–1,290 amino acids, i.e., 613 amino acids long, were used for analysis. Using I-Mutant 2.-0 and MuPro, respectively we found that out of eight nsSNPs only three amino acid substitutions (H1236Y, E1268V, and T1267I) increased protein stability and remaining (F1238L, L1253S, E1272G, V1278M, S1286) decreased the stability of the protein. In contrast, the PremPS and PoPMuSiC v3.1 predicted that all nsSNPs destabilize the protein structure **(Table 3)**.

**Table 3:**
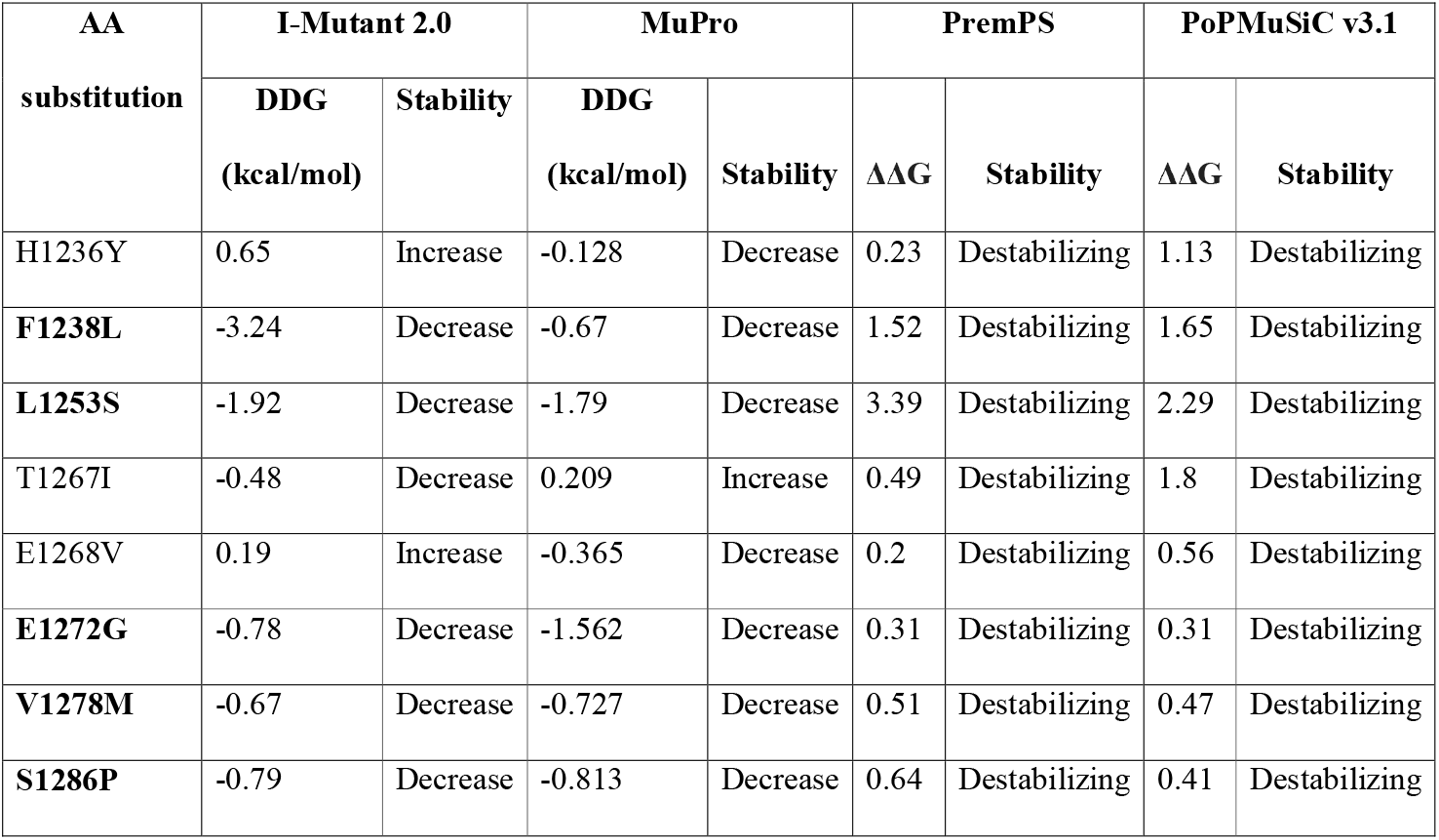
Impact of amino acid substitutions on BLM protein stability.

MutPred2 was further utilized to predict the structural, molecular, and phenotypic impacts of amino acid variations. The MutPred2 [31] webserver identified 6 out of the 8 shortlisted deleterious nsSNPs (H1236Y, F1238L, L1253S, T1267I, E1268V, and E1272G) as likely to cause structural and functional changes, with prediction scores exceeding 0.50. These changes impacted various properties, including alterations in ordered interfaces, transmembrane proteins, protein stability, metal-binding sites, and DNA-binding sites. Additionally, they led to the loss or gain of allosteric sites, catalytic and proteolytic cleavage sites, as well as alterations in post-translational modifications at different residues. (as shown in **Table 4**).

**Table 4:**
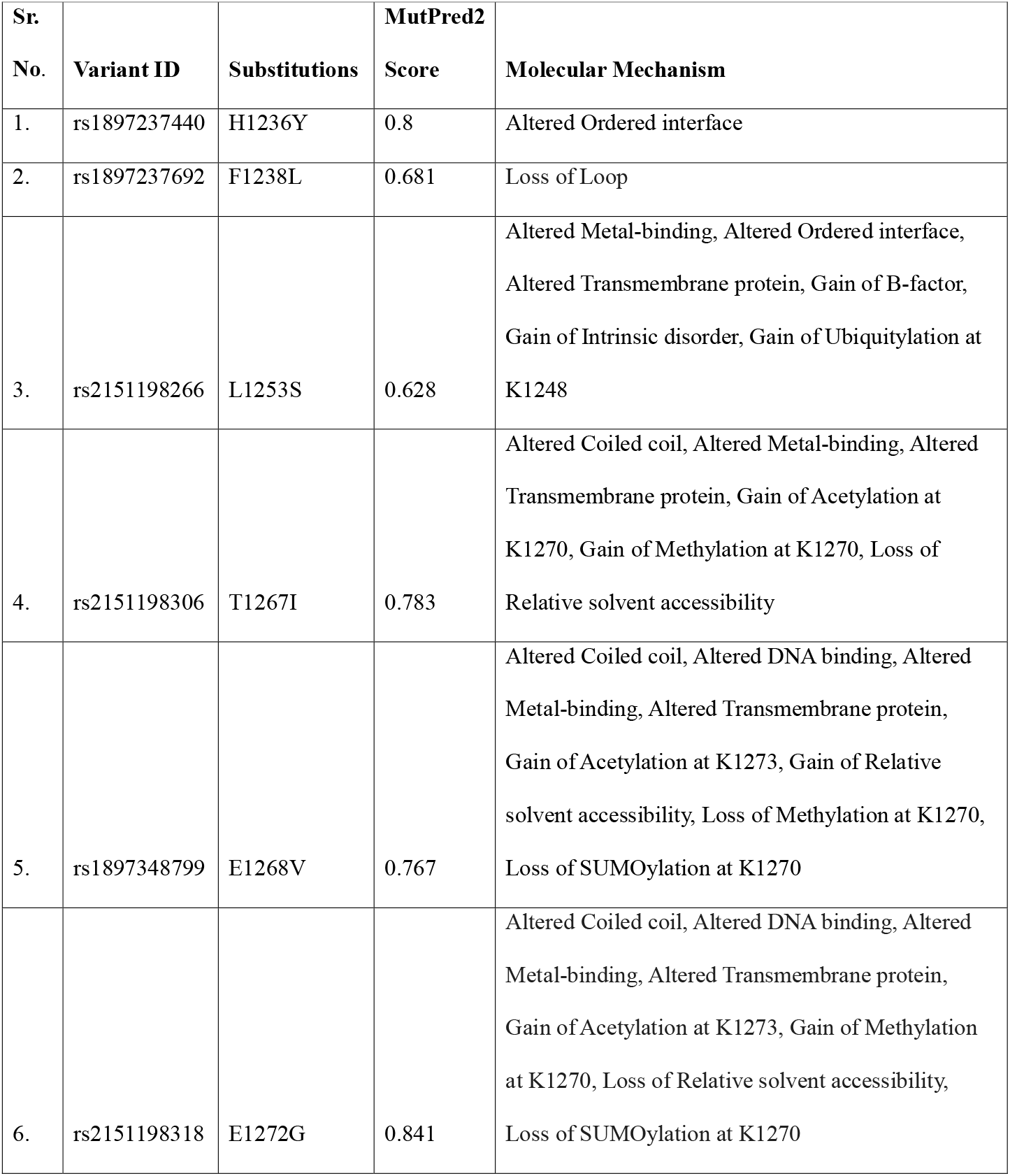
List of deleterious nsSNPs and the resulting structural and functional alterations in BLM protein.

### Association of BLM with PAPR1

STRING analysis was used to profile the protein-protein interactions involving BLM and its associated interacting proteins **Fig 4A**. Among the total 236 interactomes, PARP1 was noticed as its activity is known to be elevated in the brains of individuals with neurodegenerative diseases. We identified important associations between BLM and PARP1, an enzyme that catalyzes the addition of poly (ADP-ribose) (PAR) on target proteins. BLM and PARP1 showed similar expression profiles during cerebral cortex development **Fig 4B**. We observed both correlated co-expression and divergent expression patterns of BLM and PARP1 in normal brain tissue, underscoring their significant association, as illustrated in **Fig 4C**. Also, BLM contains a PARPylation site (1303-1333) [32] in the intrinsically disordered regions, between the HRDC and NLS sequence. By comparing the result of corGSEA between BLM and PARP1, we found differential enrichment of several pathways implicated in neurodegenerative diseases, including “Alzheimer’s Disease”, “Lu aging brain down”, “Kim all disorders duration corr down”, and “Mody Hippocampus postnatal” **Fig 4D**. This suggests the possible association of BLM and PARP1 protein in relation to neurodegenerative diseases.

**Figure 4:**
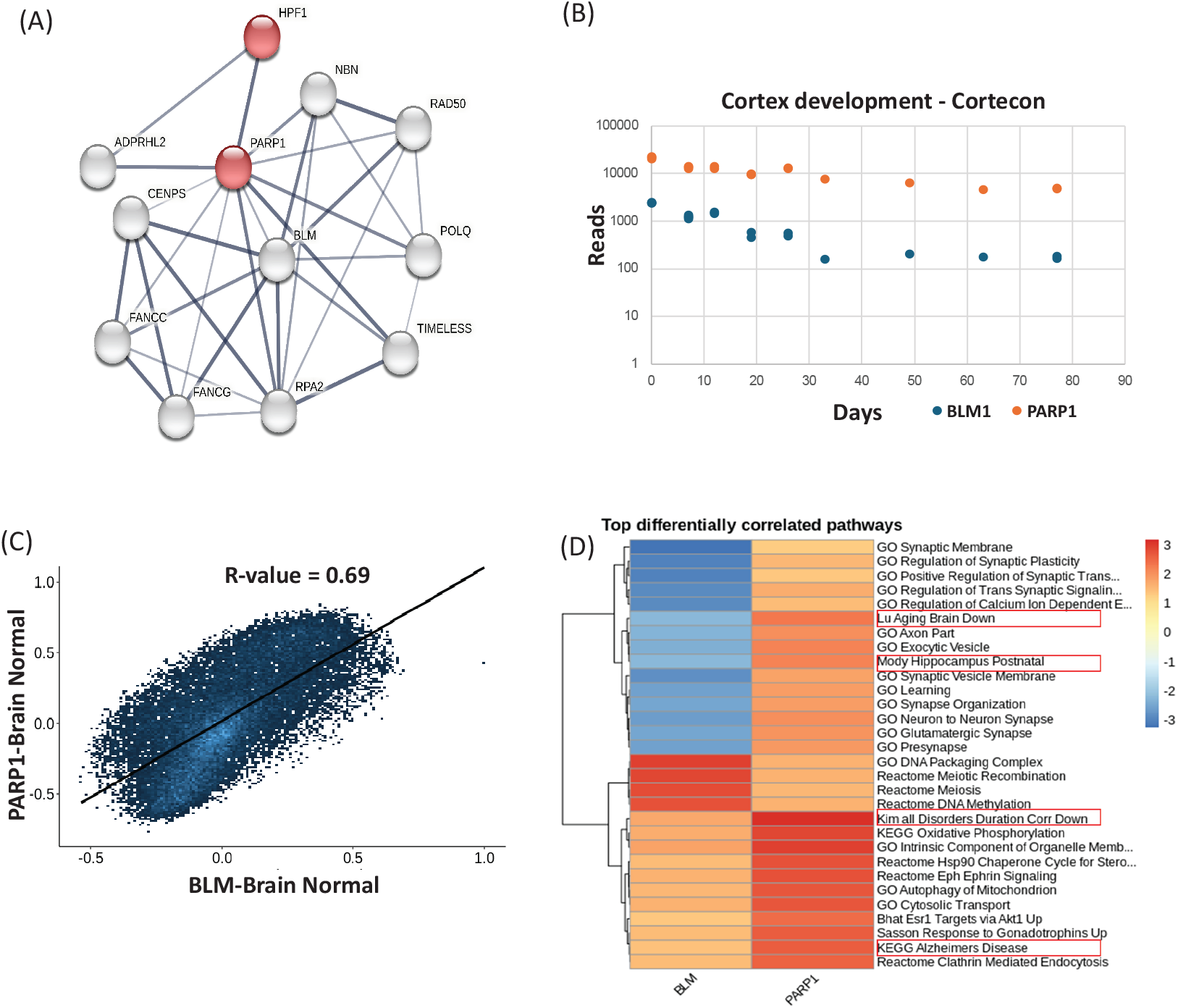
**A)** Network showing shared BLM and PARP implicated in repair mechanism. **B)** Similar pattern of BLM and PARP1 expression profiles during cortex development according to Cortecon. **D)** Scatter plot showing the relationship between genome-wide co-expression correlations for BLM and PARP1 in normal brain tissues with an R-value of 0.69 determined by Pearson correlation. **E)** Heatmap showing differential corGSEA results for BLM and PARP1 in normal tissues. The color bar displays the normalized enrichment score as determined by GSEA. Four neurodegenerative disease-related pathways are highlighted.

## Discussion

BLM belongs to the RecQ helicase family and carries an evolutionary conserved 3’ to 5’ DNA helicase activity which unwinds the double-helix DNA to produce single-stranded DNA as an intermediate required for the processes of DNA replication, repair, and recombination [33]. Immunohistochemical analysis of BLM expression across human developmental stages reveals its presence in various cell types, including Purkinje cells, neurons, thymic Hassall’s corpuscles, pancreatic beta-cells, and testicular sperm cells, with patterns varying by age and tissue. In a BS patient, BLM expression was absent in pancreatic and testicular tissues, suggesting that BLM plays a critical role in neuronal development, immune function, insulin secretion, and spermatogenesis, aligning with the major symptoms of BS [34].

Diseases associated with proteins that contain intrinsically disordered regions (IDRs) include, Neurodegenerative Diseases (Alzheimer’s Disease due to Tau protein and Amyloid-beta), Parkinson’s Disease (due to α-Synuclein), Huntington’s Disease (due to Huntingtin protein), Amyotrophic Lateral Sclerosis due to TDP-43 and FUS) [4]. These diseases are often linked to the misfolding, aggregation, or aberrant interactions of these intrinsically disordered proteins, contributing to the pathogenesis of the disease [35]. It has been shown that the proteins that have IDRs have a strong tendency to form protein aggregation, although IDR itself is not the only cause of aggregate formation [36]. Proteins such as TAR DNA-binding protein (TDP-43) in amyotrophic lateral sclerosis (ALS), α-synuclein (α-syn) in Parkinson’s disease (PD), and amyloid beta (Aβ) and tau for Alzheimer’s disease (AD) form aggregates [7]. BLM has both N-terminal and C-terminal intrinsically disordered regions [37] and HRDC domain was found to have the ability of aggregation and fibril formation similar to IDPs/IDRs proteins (**Fig 2**) (**Table 1**). Furthermore, the HRDC domain of BLM contains several non-synonymous single nucleotide polymorphisms (nsSNPs) that alter the structure or stability of the protein. Our analysis suggests various SNPs of BLM including F1238L, L1253S, and E1272G mutations might destabilize the protein based on changes in Gibbs free energy and folding energy. Additionally, these deleterious nsSNPs found in the HRDC domain may affect its structure, function, and aggregation-forming ability.

BLM interacts with PARP1, and both are involved in DNA repair mechanisms e.g. repair of double-strand break through homologous recombination and resolution of DNA replication stress. PARP1 has been identified as a BLM promoter-binding protein that negatively regulates the transcription of BLM in prostate cancer [38]. Moreover, our findings also indicate that BLM and PARP1 have a similar expression pattern in cortical development (**Fig 4**), suggesting that BLM complexes or related pathways may be modulated by PARylation. Further studies are needed to elucidate the molecular mechanisms of PARylation’s role in the intrinsically disordered region (IDR) of BLM and its impact on aggregation formation by the HRDC domain, which may have a role in neurodegenerative diseases. In brief, this is the first study that provides foundational knowledge about the HRDC domain of the BLM protein, highlighting its ability to form parallel aggregations and fibril formation. This new dimension could provide insights into its unexplored role in the pathology of neurodegenerative diseases.

## Material and Methods

The application of computational methods to gain biological insights is widely recognized and highly valued. Previous studies have demonstrated that utilizing an array of advanced tools and algorithms can significantly improve prediction accuracy. To ensure maximum precision, we employed several computational algorithms specifically designed for predicting aggregation propensity, identifying similarities to intrinsically disordered proteins (IDPs), and analyzing non-synonymous single nucleotide polymorphisms (nsSNPs) in BLM helicases, given their disease relevance. The in-silico tools utilized in this study include ProtParam, Expasy server, PASTA 2.0, PSIPRED, SIFT, PROVEAN, PolyPhen-2, SNAP2, SNP&GO, nsSNPAnalyzer, and MutPred-2. The Cortecon [39] database was used to identify the shared similarities between BLM and PARP1 expression profiling during in vitro cortex development. This comprehensive approach allowed to analyze the structural features of BLM protein and to identify disease-associated mutations in BLM genes.

### Amino Acid Sequence, Expression, Structural, and Physicochemical Properties Of BLM

The human BLM protein sequence and its other basic information were acquired from the National Center for Biotechnology Information (NCBI) (Protein ID: NP_000048.1) and the UniProt database (UniProtKB ID: P54132). We extracted RNA and protein expression data for BLM specifically in brain tissue from The Human Protein Atlas (HPA-The Human Protein Atlas) database. Comparative analysis of RNA expression (normalized transcripts per million, nTPM) and protein expression levels across various brain tissues was conducted and side-by-side bar graphs were generated to visualize the RNA and protein expression levels of BLM in these brain tissues. We used a computational tool PASTA 2.0 (Aggregation Prediction with PASTA (un6ipd.it) for the prediction of disorder and aggregation propensity in BLM protein [40]. We also used the PSIPRED and DISOPRED3 server (PSIPRED Workbench (ucl.ac.uk) to predict the secondary structure of BLM proteins based on their amino acid sequences. PSIPRED server employs neural networks and position-specific scoring matrices (PSSMs) to predict regions of the protein that are likely to form secondary structure [21]. The physicochemical parameters of the human BLM protein were calculated using the ProtParam tool available on the Expasy server (Expasy - ProtParam tool).

### Identification and Analysis of Potent nsSNPs of BLM

ENSEMBL genome browser 112 (Ensembl genome browser 112) and dbSNP database was used to retrieve nsSNPs in BLM gene using genomic coordinates “NM_000057.4” for transcript of the BLM helicase. We used the sequence-based tools such as SIFT (Sorting Intolerant From Tolerant; https://sift.bii.a-star.edu.sg/) [22], PolyPhen-2 (Polymorphism phenotyping v2; http://genetics.bwh.harvard.edu/pph2/) [23], Mutation Assessor, PhD-SNP (Predictor of human Deleterious Single Nucleotide Polymorphisms; http://snps.biofold.org/phd-snp/phd-snp), SNPs & GO (https://snps.biofold.org/snps-and-go/), Meta-SNP (http://snps.biofold.org/meta-snp/), and Screening of non-acceptable polymorphisms (SNAP) to predict the function of nonsynonymous single nucleotide polymorphism (nsSNP) [41].

SIFT (score <0.05); PolyPhen-2 (score>0.95 - probably harmful); Meta-SNP, PhD-SNP, SNPs & Go (score>0.5), PredictSNP [25] and SNAP [26] was used to determine the accuracy of the damaging effect of amino acid substitution.

### Predicting the Effect of nsSNPs on the Stability of BLM

I-Mutant 2.0 (http://folding.biofold.org/i-mutant/i-mutant2.0.html) [27] and MuPro (https://mupro.proteomics.ics.uci.edu) [28], were used to predict change in the stability of BLM upon a single nucleotide change using protein sequence as input. The associated change in free energy (ΔΔG, in kcal/mol) and stability were analyzed for the shortlisted SNPs. A negative ΔΔG value indicates a decrease in stability, while a positive ΔΔG value signifies an increase in stability. PremPS (https://lilab.jysw.suda.edu.cn/research/PremPS/) [29] and PoPMuSiC v3.1 2.1 (http://babylone.ulb.ac.be/PoPMuSiCv3.1) [30] was also used to predict change in thermodynamic stability of BLM upon a single nucleotide change. Pdb file or pdb ID was used as input file respectively by PremPS and PoPMuSiC to evaluate the effects of single mutations on protein stability by calculating the changes in unfolding and folding Gibbs free energy (ΔΔG). A positive and negative sign corresponds to destabilizing and stabilizing mutations respectively.

### Structural and Functional Changes in BLM Caused by Deleterious nsSNPs

MutPred2 (http://mutpred.mutdb.org), a popular machine learning method, is used to detect pathogenic variants as well as the structural and functional changes brought on by harmful nsSNPs [31]. It predicts their effects on protein features, enabling it to categorize them as pathogenic or benign. A list of amino acid substitutions in the relevant FASTA headers was utilized as input along with a protein sequence in FASTA format. A P-value cutoff of 0.5 and a prediction score ranging from 0.5 to 1 was employed to identify the structure and functional alterations caused by deleterious nsSNPs of BLM.

### Gene Ontology and Network Analysis of BLM-associated Proteins

BLM interactors were downloaded from BioGRID (v4.4) [12], and STRING v12.0 (the Search Tool for Retrieval of Interacting Genes and Proteins) [42] database and was used to assess its interacting genes and the protein network. Gene Ontology (GO) enrichment analysis was performed using SRPlot [19] to identify and interpret the biological functions, processes, and cellular components that are significantly associated with BLM. Finally, the gene expression correlation was analyzed using Correlation AnalyzeR [43].

